# Depletion of the protein hydration shell with increasing temperature observed by small-angle X-ray scattering and molecular simulations

**DOI:** 10.1101/2025.06.27.661897

**Authors:** Johanna-Barbara Linse, Hyun Sun Cho, Friedrich Schotte, Philip A. Anfinrud, Jochen S. Hub

## Abstract

The hydration shell is an integral part of proteins since it plays key roles for conformational transitions, molecular recognition, and enzymatic activity. While the dynamics of the hydration shell have been described by spectroscopic techniques, the structure of the hydration shell remain less understood due to the lack of hydration shell-sensitive structural probes with high spatial resolution. We combined temperature-ramp smallangle X-ray scattering (*T* -ramp SAXS) from 255–335 K with molecular simulations to show that the hydration shells of the GB3 domain and villin headpiece are remarkably temperature-sensitive. For proteins in the folded state, *T* -ramp SAXS data and explicit-solvent SAXS predictions consistently demonstrate decays of protein contrasts and radii of gyration with increasing temperature, which are shown to reflect predominantly temperature-sensitive depleting hydration shells. The depletion is not merely caused by enhanced disorder within the hydration shells but also by partial displacements of surface-coordinated water molecules. Together, *T* -ramp SAXS and explicit-solvent SAXS calculations provide a novel structural view on the protein hydration shell, which underlies temperature-dependent processes such as cold denaturation, thermophoresis, or biomolecular phase separation.

## Introduction

The hydration shell is an integral part of biomolecules since it mediates a wide range of biological functions. Water in the hydration shell of enzymes actively participates in hydrolytic enzymatic reactions, acid-base reactions, or proton transfer via the Grotthus mechanism. ^1–3^ The hydration shell furthermore orchestrates protein folding, ligand binding, or protein– protein recognition because such processes involve extensive rearrangements of protein–water and water–water interaction networks. Hydration shell water exhibits different properties as compared to bulk water. Techniques such as NMR, terahertz, sum frequency generation, or inelastic neutron scattering spectroscopy have shown that vibrational, rotational, and translational dynamics of hydration shell water are slowed down approximately two- to fivefold relative to bulk water.^4–18^

While the dynamics of the hydration shell have been extensively studied, its structural properties remain less well understood. X-ray and neutron crystallography provide insight into highly localized water molecules that coordinate with biomolecular surface residues, ^19,20^ but these techniques are largely blind to more dynamic water or to water in the second and third hydration layers. Small-angle scattering with X-rays or neutrons (SAXS/SANS) is sensitive to the hydration shell but yields data with low information content and low spatial resolution. Accordingly, SAXS/SANS has shown that many proteins exhibit hydration shells with an overall increased density relative to the bulk, which manifests in an increased radius of gyration (*R*_g_) relative to the *R*_g_ of the bare protein.^21,22^ Molecular dynamics (MD) simulations with explicit water corroborated these findings and explained the modified *R*_g_ values with a hydration shell density that is increased by *∼*6%.^23–26^ However, influences by factors such as protein geometry, surface composition, pH, or temperature on the hydration shell remain poorly characterized, largely because experimental probes of the hydration shell structure with atomic resolution are missing.

Solvation of biomolecules is governed by a delicate balance of large enthalpic and entropic contributions, which often mostly compensate to yield moderate solvation free en-ergies. Because entropic effects are amplified at high temperature, solvation is inherently temperature-dependent with implications on biomolecules and soft-matter systems. For instance, the hydrophobic effect, which drives protein folding as well as micelle and lipid membrane assembly, is temperature-dependent and entropy-driven at 22^*°*^C yet enthalpy-driven at 113^*°*^C.^27–29^ Temperature-dependent solvation drives the unfolding of proteins at low temperatures (cold denaturation)^30,31^ as well as the collapse of intrinsically disordered proteins^32^ or liquid-liquid phase separation at high temperatures.^33^ Such effects would be at odds with the naive expectation by which high temperatures would generally favor polymer disorder, thus highlighting the importance of hydration shell effects. The drift of molecules or beads along temperature gradients, an effect known as thermophoresis, thermodiffusion, or Soret effect, is driven by the temperature-dependence of the solvation entropy.^34,35^ How such effects contribute to the response of biological systems to changing temperatures, for instance during thermosensing by membrane channels or thermotaxis by bacteria, is not known. Additionally, the size and shape of detergent or polymer micelles are temperature-sensitive, which has likewise been associated with temperature-dependent solvation.^36,37^ Thus, structural insight into temperature-dependent solvation is essential to rationalize a plethora of biological or soft-matter phenomena.

We combine temperature-ramp (*T* -ramp) SAXS experiments and MD simulations to reveal how temperature controls the hydration shell structure of two proteins in their folded state, the third IgG-binding domain of Protein G (GB3) and chicken villin headpiece (also termed HP35), which have been used extensively as model proteins for biophysical studies. We developed infrastructure on the BioCARS beamline at the Advanced Light Source for acquiring SAXS data over a broad temperature range spanning 260 K to 345 K, covering supercooled conditions up to unfolded proteins. To rationalize the temperature effects on our SAXS data by atomic means, we use all-atom MD simulations combined with explicit-solvent SAXS calculations.^25,38^ Our strategy is supported by recent findings that the overall hydration shell density obtained by MD simulations with many protein and water force fields aligns accurately with consensus SAXS/SANS data.^39,40^ Our *T* -ramp SAXS data and MD simulations are quantitatively consistent and demonstrate that —in the folded state— the protein hydration shell is remarkably temperature-sensitive as shown by partial displacements of surface-coordinated water molecules.

## Results

Figure 1A (upper row) presents the three-dimensional electron density of solvent around the GB3 domain, computed from an MD simulation with the TIP4P/2005 water model, which has been parametrized to reproduce water properties across a broad temperature range. ^41^ By using position restraints on all heavy atoms, conformational fluctuation of solvent-exposed protein atoms were excluded, thereby yielding a spatially well-defined hydration shell with a first and a second hydration layer (orange and blue densities, respectively). The density maps reveal that increasing the temperature from 250 K to 350 K leads to a partial loss of hydration shell structure, as evident from decreased densities of the first and second hydration layer. These1‘124z simulations provide the atomistic rationale for interpreting temperature-dependent scattering intensities observed in SAXS experiments.

**Figure 1:**
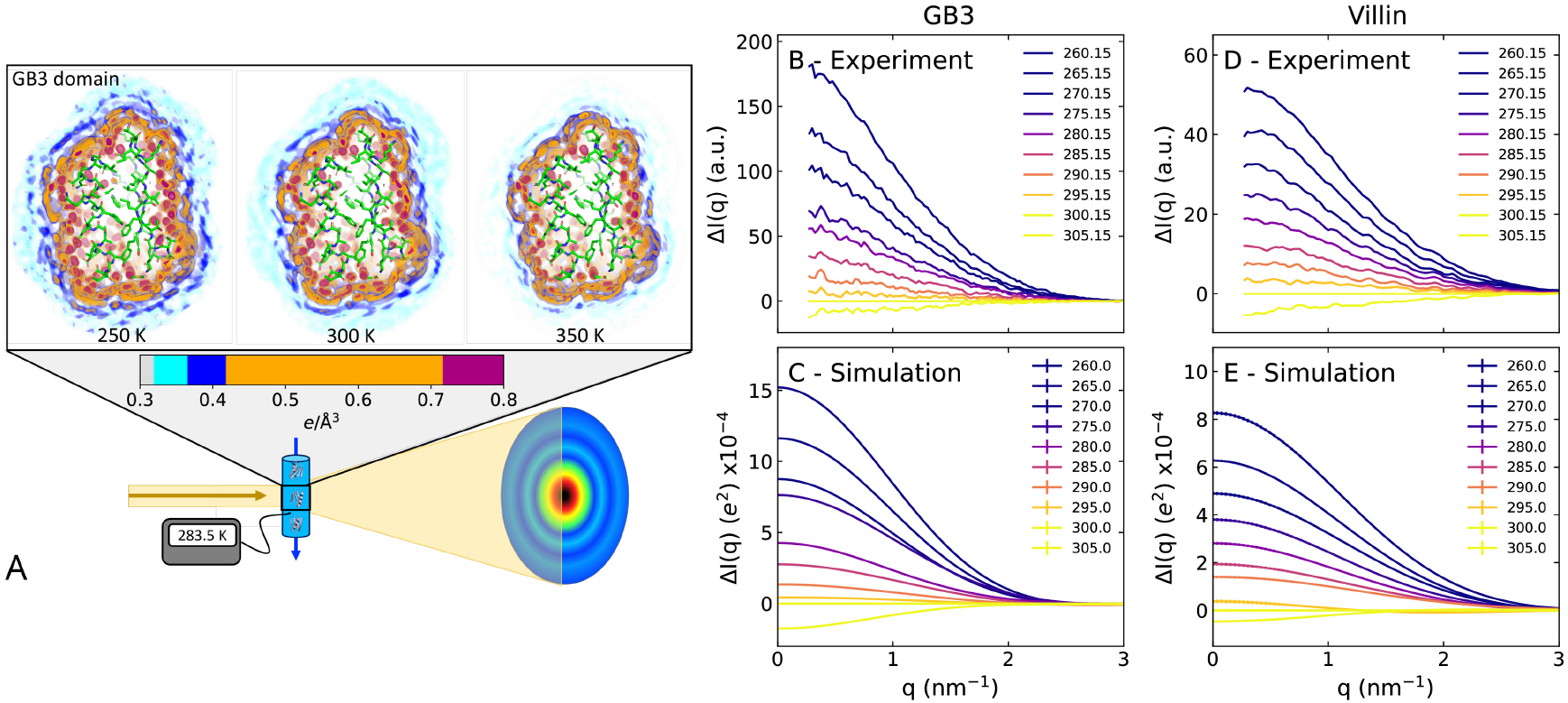
(A) Illustration of temperature-ramp SAXS experiments. Upper row: threedimensional electron density around the GB3 domain at 250 K, 300 K and 350 K from MD simulations with restrained heavy atoms. The protein is shown as sticks, the solvent electron density in colors from cyan to purple, see color bar, revealing the depletion of the hydration shell structure with increasing temperature. (B) Scattering intensity difference relative to 300 K, Δ*I*(*q*), of the GB3 domain at temperatures spanning *∼*260 K to *∼*305 K (see legend) from experiments or (C) from simulations. (D) Δ*I*(*q*) for the villin headpiece from experiments or (E) from simulations. Absolute SAXS curves are shown in Figs. S1 and S2.

SAXS data for the GB3 domain and villin headpiece were acquired on the BioCARS beamline at the Advance Light Source using a *T* -ramp protocol that generates scattering images while repeatedly ramping the sample temperature between 260 K and up to 393 K (illustrated in Fig. 1A, lower row). The SAXS curves of both proteins exhibited clear temperature dependence (Figs. S1A and S2A). Thanks in part to the small volume of protein solution in the HF-etched, temperature-controlled capillary, the solution could be repeatably supercooled to 260 K without freezing. The temperature dependence of the SAXS curves is highlighted by the difference intensities Δ*I*(*q*) relative to the experimental reference temperature of 300.15 K (Fig. 1B/D). In this study, we focus on two key parameters that are frequently obtained from the small-angle regime: (i) the forward scattering intensity *I*_0_ = *I*(*q* = 0), which is given by the square of the contrast between solute and solvent (in number of electrons), scaled in experiment by a concentration-dependent factor; and (ii) the radius of gyration *R*_g_. These parameters were obtained via Guinier analysis, 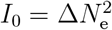, where *I*(*q*) is the SAXS curve and *q* the momentum transfer.

Our SAXS experiments revealed that the forward scattering *I*_0_ decreased by more than 20% for both the GB3 domain and villin headpiece across the measured temperature range, demonstrating a decreasing electron density contrast that plateaued at *∼*320 K for the GB3 domain and at *∼*335 K for the villin headpiece, respectively (Fig. 2A/D, black dots). The *R*_g_ values decreased by 0.4 Å for the GB3 domain and by 0.3 Å for villin headpiece between 257.15 K and *∼*300 K (Fig. 3A/C, black dots). This trend is followed by a slight increase of *R*_g_ up to *∼*330 K, likely indicating enhanced conformational fluctuations, followed by sharp *R*_g_ increases indicating protein unfolding. The temperatures of sharp *R*_g_ increase are compatible with previously reported melting temperatures of the GB3 domain and villin headpiece.^42–44^

**Figure 2:**
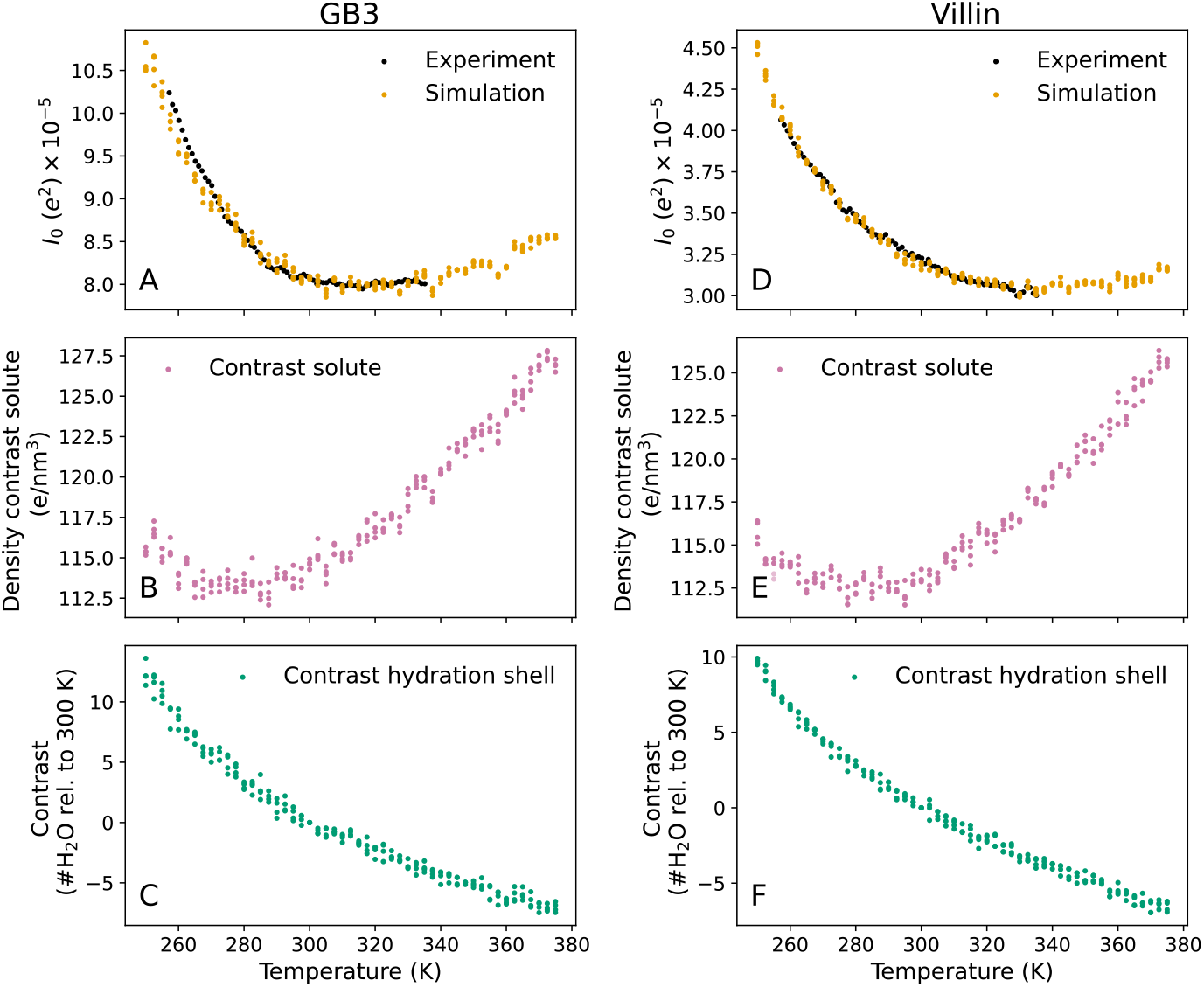
Forward scattering *I*_0_ and decomposition of the solute–solvent contrast into contributions from the bare protein and the hydration shell for (A–C) the GB3 domain and (D–F) villin headpiece. (A/D) *I*_0_ versus temperature from experiment (black) and backbone-restrained MD simulations with explicit-solvent SAXS calculations (orange). The experimental curves were scaled by a constant factor to the simulation data in the temperature range below 303 K. (B/E) Density contrast of the backbone-restrained bare protein from MD simulations reflecting the temperature dependence of the solvent density. (C/F) Contrast of the hydration shell in number of water molecules relative to 300 K. In all panels, colored dots indicate simulation results from four independent simulations per temperature. The analysis of panels A–F visualized as contrasts in number of electrons is shown in Fig. S4.

**Figure 3:**
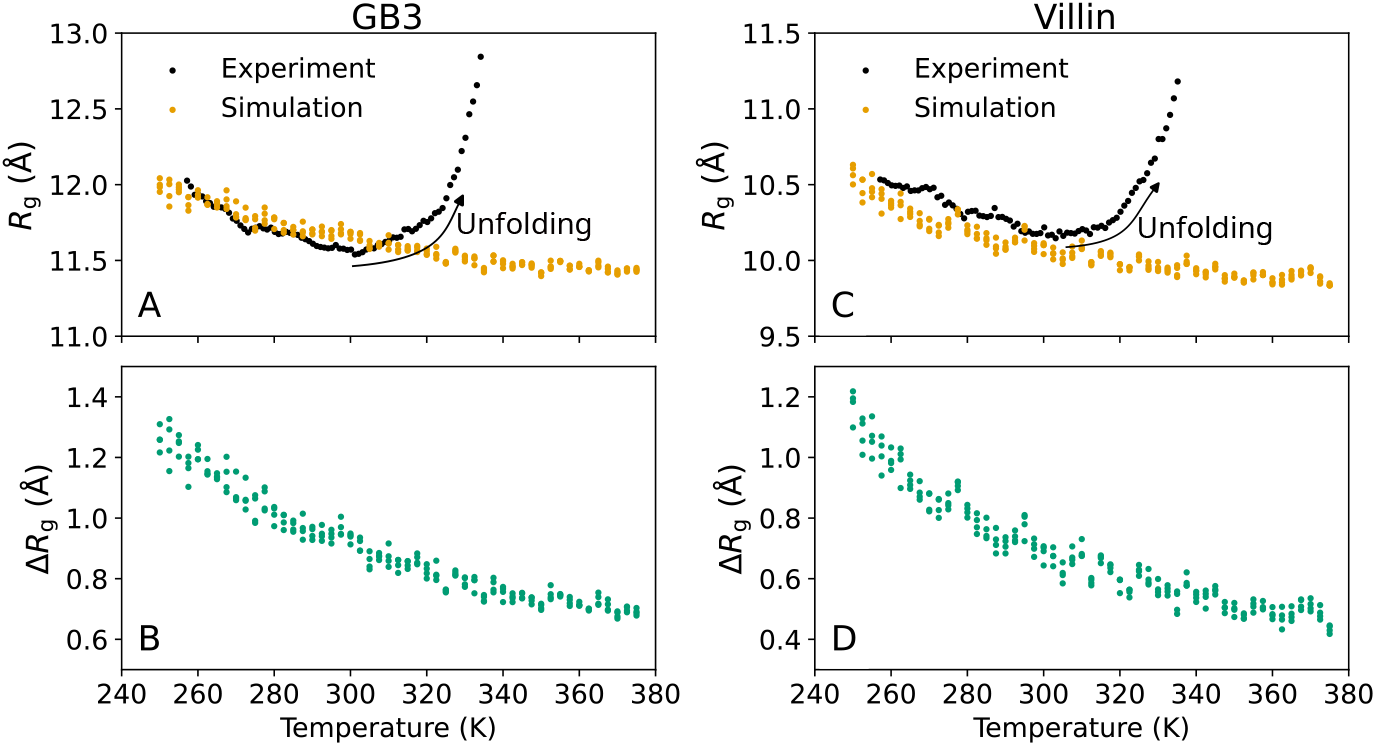
Radius of gyration *R*_g_ versus temperature for (A/B) the GB3 domain and (C/D) villin headpiece. (A/C) *R*_g_ from Guinier analysis from experiment (black) or backbone-restrained MD simulations (orange). (B/D) From MD simulations and explicit-solvent SAXS calculations, difference Δ*R*_g_ between *R*_g_ from Guinier analysis (including hydration shell contributions) and *R*_g_ of the bare protein. Decrease of *R*_g_ with temperature in backbonerestrained simulations demonstrates the depletion of the hydration shell. Colored dots indicate simulation results from four independent simulations per temperature.

We hypothesized that the decreases in *I*_0_ and *R*_g_ between 260 K and 300 K, where the proteins are folded, reflect gradual depletions of the protein hydration shells. To test this, we carried out MD simulations of the GB3 domain and villin headpiece (Fig. S3) over a temperature range of 250 K to 375 K in steps of 2.5 K. SAXS curves were computed from MD simulations according to the WAXSiS method, thereby taking the explicit solvent into account.^25,45^ To isolate the effect of the solvent and to prevent unfolding, the simulations were carried out with position restraints on backbone atoms; thus, variations of the computed SAXS curve were purely caused by temperature-dependent variations of hydration shell and excluded solvent densities.

In line with our SAXS experiments, the computed SAXS curves were temperature-dependent, as shown by the absolute scattering curves (Figs. S1B, S2B) and by difference curves relative to 300 K (Fig. 1C/E). The *I*_0_ and *R*_g_ values taken from the calculated curves via Guinier analysis are in good agreement with the experimental values in the *≤* 300 K regime, where the proteins remain fully folded. The agreement is remarkable considering that, in our explicit-solvent SAXS calculations, neither the hydration shell nor the excluded solvent density were fitted against the experimental data. The agreement implies that (i) thermal expansion of the protein, which is excluded in our MD simulations with backbone restraints, has only a small effect on *I*_0_ and *R*_g_ in this temperature range; and (ii) that the MD simulations with the TIP4P/2005 water model provide a realistic representation of the hydration shell structure across a broad temperature range.

To isolate the effect of temperature on the hydration shell, we decomposed the total contrast Δ*N*_e_ (in number of electrons) into contributions from the protein and the hydration shell. The forward scattering follows 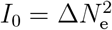 and the total contrast is

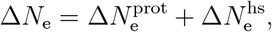

where 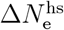 is the contrast imposed by the hydration shell and 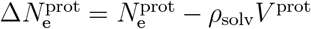 the contrast of the bare protein with the temperature-dependent solvent density *ρ*_solv_, *V* ^prot^ the protein volume, and 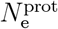 the number of electrons in the protein. Figures 2B/E show the electron density contrast of the bare proteins, 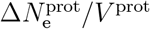, reflecting purely the temperature dependence of the water density as the protein volumes are nearly fixed. Here, the convex shapes of the density contrast curves reflect the well-known maximum of the water density at 4^*°*^C, with the density decreasing at both low and high temperatures. Critically, the contrast between the hydration shell and bulk water 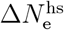 greatly decreases with increasing temperature, demonstrating a depleting hydration shell structure (Fig. 2C/F). Specifically, relative to 300 K, the hydration shell of both the GB3 domain and villin headpiece contain approximately 10 additional water molecules at 250 K and 5 fewer water molecules at 360 K (Fig. 2C/F). Over the entire temperature range simulated here, the hydration shells of the GB3 domain and villin headpiece lose 19 and 17 water molecules, respectively. We note that the quantitative decomposition of the Δ*N*_e_ into contributions from protein and hydration shell depends on the definition of the protein volume and, thus, a different definition may lead to a constant shift between 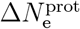 and 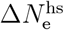,^26^ which would, however, not alter the trends as function of temperature. Together, this analysis supports our hypothesis that the experimentally and computationally observed decay of *I*_0_ reflects a temperature-sensitive depletion of the hydration shell.

As a second indicator of the hydration shell, we computed from the simulations the increase of the radius of gyration owing to the hydration shell, 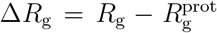, defined as the *R*_g_ from Guinier analysis relative to the 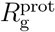of the bare protein (Fig. 3B/D). Recently, we found excellent agreement for Δ*R*_g_ values at room temperature obtained from consensus SAXS/SANS data and from MD simulations using various protein force fields and water models, including the TIP4P/2005 water model employed here.^39,40^ According to the simulations, Δ*R*_g_ decays with increasing temperature by *∼*0.6 Å, reflecting the diminishing hydration shell in line with the decaying *I*_0_ discussed above. Considering that (i) 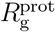was fixed in simulations due to backbone restraints, and (ii) *R*_g_ values from simulations agreed with experimental values in the *T ≤* 300 K regime where the proteins are folded (Fig. 3A/C), this analysis suggests that the *R*_g_ decays in experiments up to 300 K are likewise signatures of depleting hydration shells with increasing temperature.

Having established that the temperature-dependent hydration shell contrast agrees between simulation and experiment, we used the simulations to gain structural and energetic insights into the hydration shell and its interaction with the protein. Visual inspection of the hydration shell densities confirms that the hydration shells are gradually depleted with increasing temperature (Movies S1, S2; Fig. 1A). To resolve contributions from individual surface-bound water molecules, we computed the density differences relative to 300 K from an additional series of simulations with restraints on all heavy atoms, thereby suppressing side chains fluctuations and leading to a spatially well-resolved hydration shell structure (Fig. 4, Movies S3, S4). In addition to the overall depletion of the first and second hydration layers, these density difference maps reveal the loss of numerous localized solvent densities. Visual inspection showed that most of these localized density differences originate from molecules that are coordinated via hydrogen bonds with the protein, whereas few density differences originate from water molecules trapped in small hydrophobic pockets (Fig. 4, red and blue spots). Critically, localized densities from surface-bound water at low temperatures are not merely smeared out at higher temperatures but partially lost. This finding is supported by the hydration shell densities as function of distance from the Van-der-Waals surface of the proteins, which reveal a partial loss of the first and second hydration shell densities peaks (Figs. S5A, S6A). These structural changes are accompanied by a *∼*13% decrease in both protein–water interaction energies and the number of hydrogen bonds over the simulated temperature range (Figs. S5B/C, S6B/C). Nevertheless, even at 350 K, the overall hydration shell structure with its pronounced first and shallow second peak remains intact (Figs. S5A, S6A), suggesting that the decrease of *I*_0_ and *R*_g_ reflect a partial but not complete loss of the protein–water coordination with increasing temperature.

**Figure 4:**
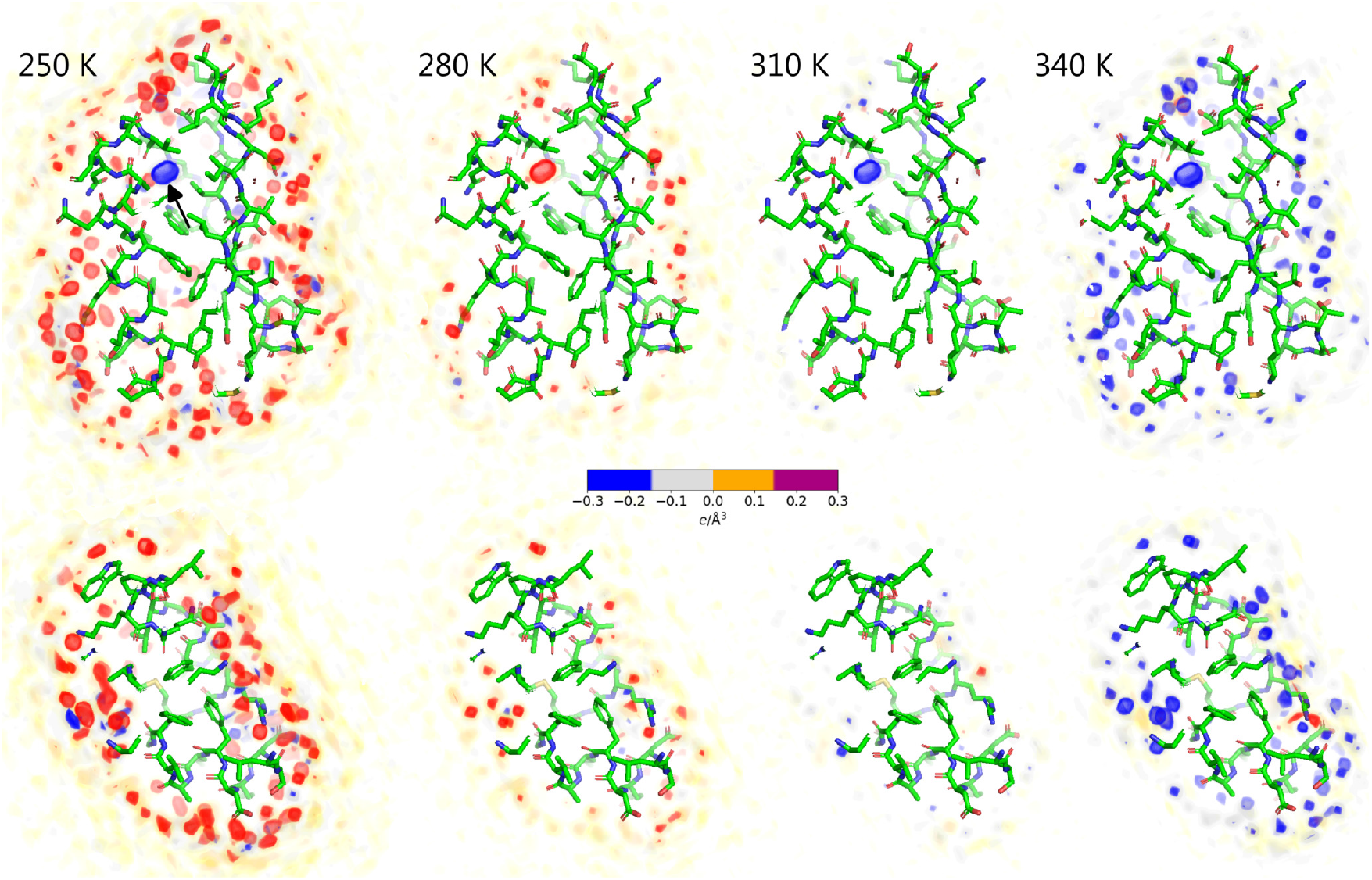
Solvent density difference relative to 300 K for the GB3 domain (upper row) and villin headpiece (lower row) for temperatures of (from left to right) 250 K, 280 K, 310 K, or 340 K. Densities were computed from MD simulations with restraints on all heavy protein atoms. Increased and decreased densities relative to 300 K are shown as red and blue densities, respectively (see color bar), revealing the depletion of surface-bound densities with increasing temperature. The localized density (black arrow) refers to a randomly placed water molecule within a GB3 cavity, which is not exchanged with the bulk within simulation time. Proteins are shown as sticks.

In addition to analyzing protein–water coordination discussed above, we investigated how temperature affects the internal structure of the hydration shell. To this end, we computed the number of water–water hydrogen bonds within a distance of 9 Å from the protein surfaces of the GB3 domain and villin headpiece (Figs. S5D, S6D), revealing a *∼*20% decrease over the simulated temperature range. This trend, together with increasingly smeared out water–water radial distribution function within the hydration shell (Fig. S7), demonstrates a considerable loss of internal water structure within the hydration shell, consistent with previous computational and spectroscopic studies.^46–49^ At high temperatures, the decrease in the number of hydrogen bonds is more pronounced in the hydration shell as compared to the bulk solvent, suggesting that the hydration shell is more temperature-sensitive than bulk water (Figs. S5D, S6D). Together, these analyses suggest that increasing temperature leads not only to a loss of coordinated water densities at the protein surface, but furthermore to a generally less structured hydration shell, as demonstrated by fewer water–protein and water–water hydrogen bonds, more dispersed protein–water and water–water correlations, and reduced enthalpic protein–water interactions. These findings provide the structural rationale for the decreasing contrast and Δ*R*_g_ values observed by our SAXS experiments and explicit-solvent SAXS calculations.

In summary, using a novel *T* -ramp SAXS setup at BioCARS, we investigated the GB3 domain and villin headpiece across a broad temperature range, spanning supercooled conditions to protein unfolding. At lower temperatures, where the proteins remain folded, the SAXS data revealed systematic decreases in the protein’s electron density contrasts and radii of gyration. MD simulations combined with explicit-solvent SAXS calculations showed excellent agreement with the experimental data and attributed these trends primarily to the temperature-dependent depletion of the hydration shell. The depletion is not solely caused by increased water disorder, as is expected at increasing temperatures, but involves partial displacement of surface-coordinated water molecules. Together, the SAXS and simulations provide detailed structural insight into the protein hydration shell, highlighting its remarkable sensitivity to temperature, as underlying temperature-sensitive biological processes such as cold denaturation, thermophoresis, and biomolecular phase separation.

## Methods

### *T* -ramp SAXS experiments

The 35-residue villin headpiece subdomain (LSDED FKAVF GMTRS AFANL PLWKQ QHLKK EKGLF) was obtained from California Peptide Research Inc. The peptide was dialyzed against 20 mM acetate buffer at pH 4.9 with 150 mM NaCl. The GB3 domain (MQYKL VINGK TLKGE TTTKA VDAET AEKAF KQYAN DNGVD GVWTY DDATK TFTVTE) was expressed and purified as described previously^50^ and dissolved in 40 mM acetate buffer with 150mM NaCl, 0.05% NaN3, 5 mM DTT at pH 5.5.

Temperature-dependent small- and wide-angle X-ray scattering (SAXS-WAXS) data were acquired on the BioCARS 14IDB beamline at the Advanced Photon Source.^51–54^ Briefly, a peristaltic pump circulates in a closed loop sample solution through a 560-mm long capillary (Polymicro TSP250350) that is supported on a home-built high-speed XYZ stage. To minimize scattering from the capillary walls, the region where X-rays pass through was HF-etched to a wall thickness of approximately 15–20 µm. X-rays passing through the capillary are scattered and detected on a large area Rayonix MS340-HS detector positioned 186 mm downstream from the capillary. Thanks to the small, 0.51-mm diameter beamstop located near the midpoint between the sample and detector, the range of *q* accessible with 12 keV photons spans from 0.02 to 5.2 Å^*™*1^. The high-speed stage translates the sample capillary at constant velocity over a *∼*20-mm range during which 40 X-ray shots are transmitted through the sample at *∼*40 Hz with a separation of 0.5 mm along the capillary, thereby distributing the X-ray dosage over a large volume of sample. During the return stroke, the X-ray scattering image is saved and a fresh aliquot of solution is drawn from a *∼*120 µL sample reservoir and pushed into the capillary. A home-made temperature controller ramped the sample temperature up and down between *™*16 to 120^*°*^C at a rate of nearly 1^*°*^C/sec, repeating the ramp three times for each dataset. Scattering data were acquired at each of three different concentrations (2.2, 6.9, 20 mg/ml for villin headpiece and 2.7, 8.4, 24.7 mg/ml for GB3). To prevent boiling when ramping the capillary temperature to 120^*°*^C, the sample reservoir was pressured with helium at 3 atm. They were averaged, extrapolated to the infinite dilution limit, and Guinier analyzed to generate temperature-dependent *I*_0_ and *R*_g_ curves.

### Simulation setup and explicit-solvent SAXS calculations

Structures of the villin headpiece and the GB3 domain were taken from the protein data bank (PDB; codes 1yrf^55^ and 2oed,^56^ respectively). Crystal water was kept in the structures and hydrogen atoms were added with the pdb2gmx software.^57^ MD simulations were carried out with the GROMACS software, version 2021.7.^57^ Interactions of the proteins were described with the amber99SB-ildn force field (ff99SB-ildn). ^58^ The starting structures were placed in a dodecahedral box, where the distance between the protein and the box edges was at least nm, and solvated in TIP4P/2005^41^ water. The charges of the proteins were neutralized by adding Na^+^ or Cl^*™*^ ions. After 400 steps of minimization with the steepest decent algorithm, the systems were equilibrated for 100 ps with harmonic position restraints applied to the heavy atoms of the proteins (force constant 2000 kJ mol^*™*1^nm^*™*2^). Subsequently, the production runs were carried out for 50 ns with harmonic position restraints applied to backbone atoms (force constant 2000 kJ mol^*™*1^nm^*™*2^) at temperatures between 250 K and 375 K in steps of 2.5 K. For each protein and temperature, four independent simulations were carried out. The equations of motion were integrated using a leap-frog algorithm.^59^ The temperature was controlled using velocity rescaling (*τ* =1 ps). ^60^ The pressure was controlled at 1 bar with the Berendsen thermostat (*τ* =1 ps)^61^ and with the Parrinello-Rahman thermostat (*τ* =5 ps)^62^ during equilibration and productions simulation, respectively. The geometry of the water molecules was constrained with the SETTLE algorithm,^63^ and LINCS^64^ was used to constrain all other bond length. A time step of 2 fs was used. Dispersive interactions and short-range repulsion were described by a Lennard-Jones potential, which was cutoff at 1 nm. The pressure and energy were corrected for missing dispersion corrections beyond the cut-off. Neighbor lists were updated with the Verlet scheme. Coulomb interactions were computed with the smooth particle-mesh Ewald method.^65,66^ We used a Fourier spacing of approximately 0.12 nm, which was optimized by the GROMACS mdrun module at the beginning of each simulation.

To compute the SAXS curves, 2500 simulation frames were used from the time interval between 0 and 50 ns. The SAXS calculations were performed with GROMACS-SWAXS, as also implemented by our webserver WAXSiS.^25,45,67^ The implementation, documentation, and tutorials are available at https://gitlab.com/cbjh/gromacs-swaxs. A spatial envelope was built around the protein atoms from all frames. The distance between the protein and the envelope surface was at least 12 Å, such that all water atoms of the hydration shell were within the envelope. Solvent atoms inside the envelope contributed to the calculated SAXS curves. The buffer subtraction was carried out using 2498 simulation frames of a pure-water simulation box, which was simulated for 50 ns at the same temperature as the protein simulation and large enough to enclose the envelope. The orientational average was carried out using 150 **q**-vectors for each absolute value of **q**. The solvent electron density was corrected to the temperature-dependent experimental value of the respective temperature such as 334 e/nm^3^ for 298.15 K, as described previously.^67^ No fitting parameters owing to the hydration layer or excluded solvent were used, implying that the *R*_g_ values were not adjusted by fitting parameters.

Three-dimensional solvent densities around the GB3 domain (Figs. 1A and 4, Movies S1– S4) were computed with the mdrun module of GROMACS-SWAXS, using the environment variable GMX_WAXS_GRID_DENSITY, and visualized with PyMOL.^68^ One-dimensional solvent densities around the protein (Figs. S5A, S6A) were computed with the gmx genenv module of GROMACS-SWAXS. Here, three- and one-dimensional densities were computed from simulation with restrained heavy atoms, which prevents smearing out of solvent densities near the protein surface due to side-chain fluctuations. Protein-water interaction energies (Figs. S5B, S6B) were computed as the sum of protein–water Lennard-Jones and short-range Coulomb interactions. Hydrogen bonds were computed with the gmx hbond module using standard settings.

RDFs between water oxygen atoms within the hydration shell were computed with an inhouse modification of gmx genenv using simulations with restraints on all backbone atoms. To this end, envelopes were constructed at distances of 0 Å, 3 Å, 5 Å, or 7 Å from the Vander-Waals surface of the protein. For each MD frame of the production simulations, water oxygen atoms between (i) the envelope at 0Å distance and (ii) the envelope at *x* Å distance were selected, where *x* ∈ {3, 5, 7}. Thereby, oxygen atoms within a distance of *x* Å from the protein were selected.

To decompose the total contrast into the contrast of the bare protein and the hydration shell, we defined the protein volume *V*_prot_ as the cavity volume calculated with the 3V volume calculator^69^ with a grid size of 0.1 Å and a probe radius of 1.4 Å, as used previously.^70,71^ The number of solute electrons 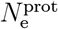was taken from the atomic form factors at zero scattering angle, as defined by the Cromer-Mann parameters of the respective atoms.

## Acknowledgement

JBL and JSH were supported by the Deutsche Forschungsgemeinschaft (DFG, German Research Foundation) via grants HU 1971/3-1 and INST 256/539-1. The temperature-dependent SAXS studies were performed on the BioCARS 14IDB beamline at the Advanced Photon Source, a U.S. Department of Energy (DOE) Office of Science user facility operated for the DOE Office of Science by Argonne National Laboratory under Contract No. DE-AC02-06CH11357. Use of BioCARS was supported by the National Institute of General Medical Sciences of the National Institutes of Health under grant number P41 GM118217. This research was supported by the Intramural Research Program of the National Institute of Diabetes and Digestive and Kidney Diseases, National Institutes of Health.

## Supporting Information Available

The Supporting Information is available free of charge on the ACS Publications website.

- Figures S1–S7: absolute SAXS curves, simulation systems, solvent density correlations, hydrogen bonds, interaction energies. Movies S1–S4: three-dimensional density maps for all temperatures.

## Code availability statement

A Gromacs variant GROMACS-SWAXS that implements explicit-solvent SAXS calculations is freely available at https://gitlab.com/cbjh/gromacs-swaxs. Documentation and tutorials are available at https://cbjh.gitlab.io/gromacs-swaxs-docs/.

## TOC Graphic

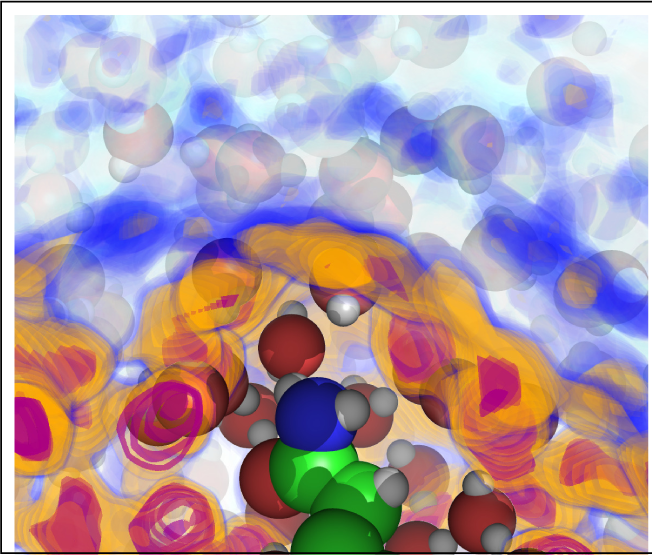

## Notes

### Competing Interest Statement

The authors have declared no competing interest.

