## Supplementary PDF for "Depletion of the protein hydration shell with increasing temperature observed by small-angle X-ray scattering and molecular simulations"

*§Correspnding author*

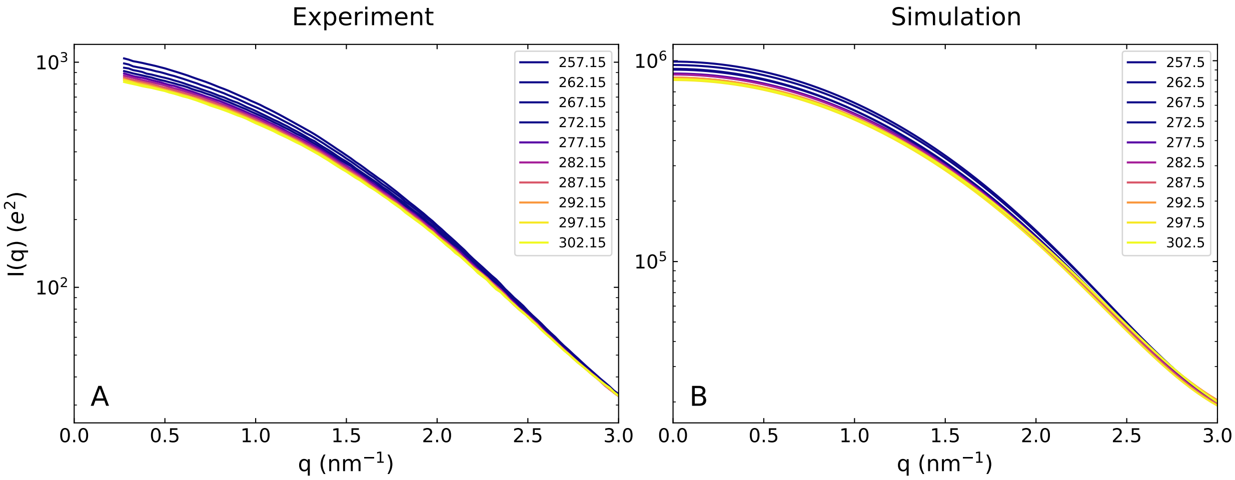


Figure S1: SAXS curves of the GB3 domain from (A) experiment and (B) MD simulation over the temperature range shown in the figure legend.

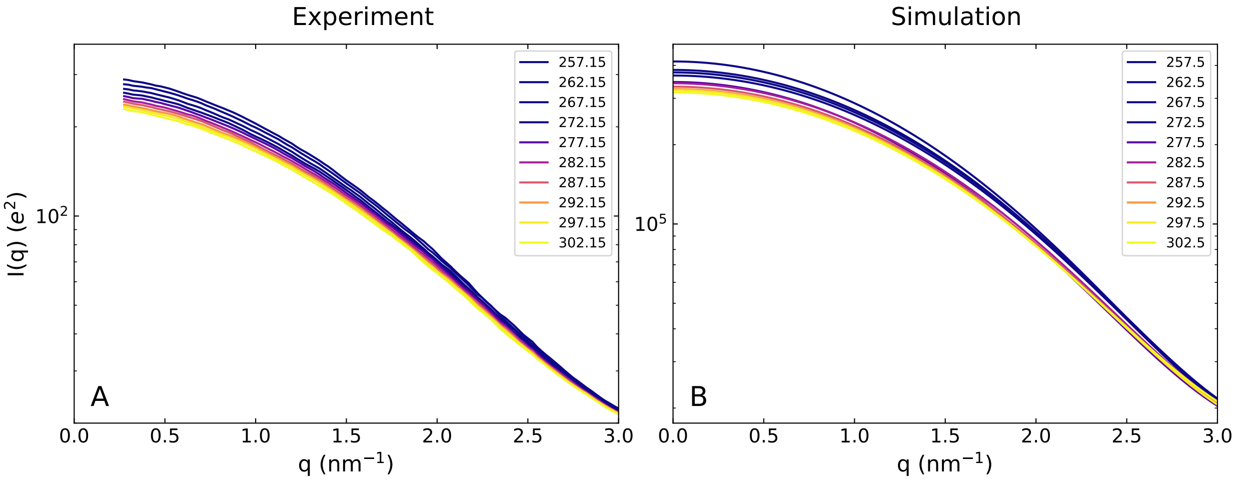


Figure S2: SAXS curves of villin head headpiece from (A) experiment and (B) MD simulation over the temperature range shown in the figure legend.

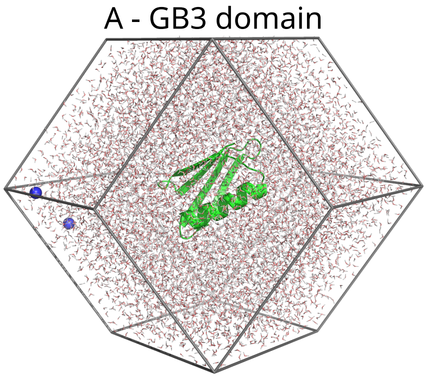


Figure S3: Simulation systems of (A) the GB3 domain and (B) villin headpiece in dodecaheral simulation boxes. Proteins are shown in green cartoon representation, water as red/white sticks, and counter ions as blue spheres.

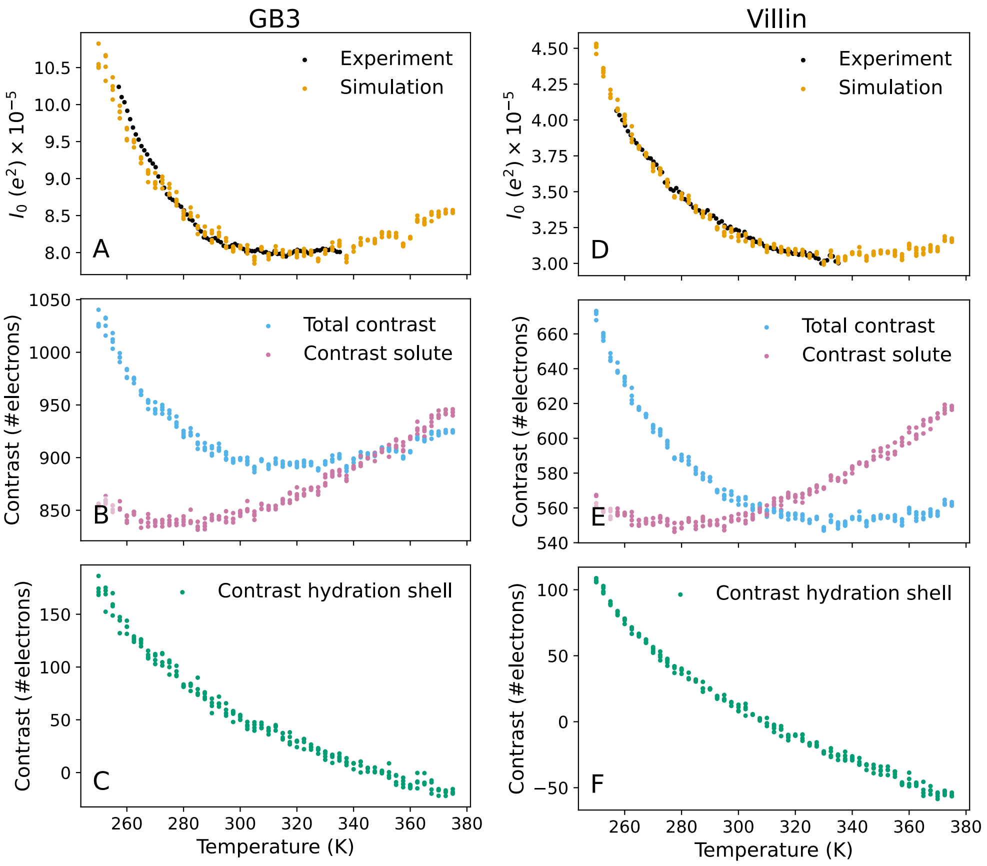


Figure S4: Forward scattering $I_0$ and analysis of contrast in number of electrons versus temperature for (A–C) the GB3 domain and (D–F) villin headpiece. Same data as presented in Fig. 2, however presented to directly illustrate our analysis. (A/D) $I_0$ from experiment (black) and backbone-restrained MD simulations (orange) versus temperature. The experimental data was scaled by a constant factor to the simulation data in the temperature range below 303 K. (B/E) Total contrast in number of electrons computed as $I_0^{1/2}$ from MD simulation (blue) and contrast $\Delta N_\mathrm{e}^\mathrm{prot}$ due to the solute (pink). (C/F) Contrast of the hydration shell $\Delta N_\mathrm{e}^\mathrm{hs}$ given as the difference between the total contrast (B/E, blue) and solute contrast (B/E, pink). Colored dots in panels (A–F) indicate simulation results from four independent simulation replicates per temperature.

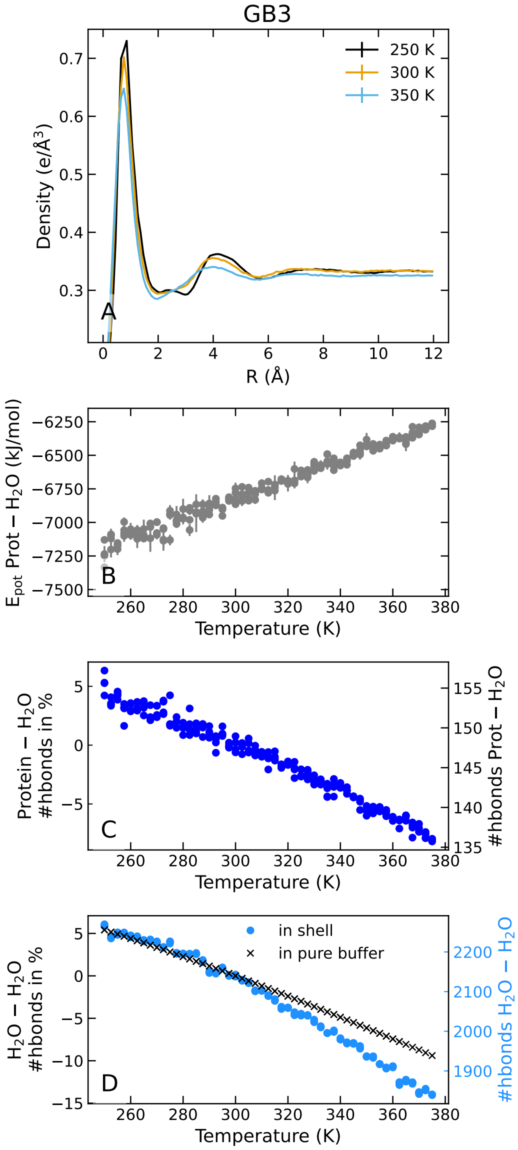


Figure S5: Analysis of the hydration shell of the GB3 domain. (A) Solvent density versus distance from the Van-der-Waals surface of the protein as averaged over the protein surface at three temperatures (see legend). (B) Protein–water interaction energy versus temperature computed as sum of Lennard-Jones and short-range Coulomb energies. (C) Number of protein–water hydrogen bonds, plotted either as total number of hydrogen bonds (right ordinate) or as change of number of hydrogen bonds relative to 300 K (left ordinate). (D) Number of water–water hydrogen bonds in the hydration shell (blue dots), defines at water within a distance of 9 Å from the protein surface. Plotted as number of hydrogen bonds (right ordinate) and relative to 300 K (left ordinate). For reference, the relative change of the number of water–water hydrogen bonds is shown for bulk water (black crosses, left black ordinate). Dots in panels B/C indicate simulation results from four independent simulation replicates per temperature.

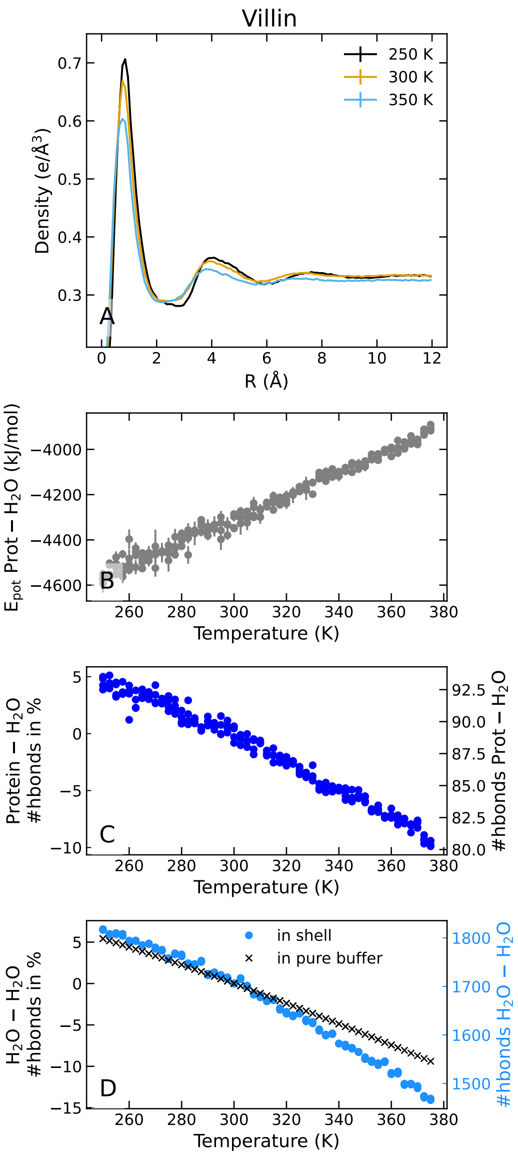


Figure S6: Analysis of the hydration shell of villin. (A) Solvent density versus distance from the Van-der-Waals surface of the protein as averaged over the protein surface at three temperatures (see legend). (B) Protein–water interaction energy versus temperature computed as sum of Lennard-Jones and short-range Coulomb energies. (C) Number of protein–water hydrogen bonds, plotted either as total number of hydrogen bonds (right ordinate) or as change of number of hydrogen bonds relative to 300 K (left ordinate). (D) Number of water–water hydrogen bonds in the hydration shell (blue dots), defines at water within a distance of 9 Å from the protein surface. Plotted as number of hydrogen bonds (right ordinate) and relative to 300 K (left ordinate). For reference, the relative change of the number of water–water hydrogen bonds is shown for bulk water (black crosses, left black ordinate). Dots in panels B/C indicate simulation results from four independent simulation replicates per temperature.

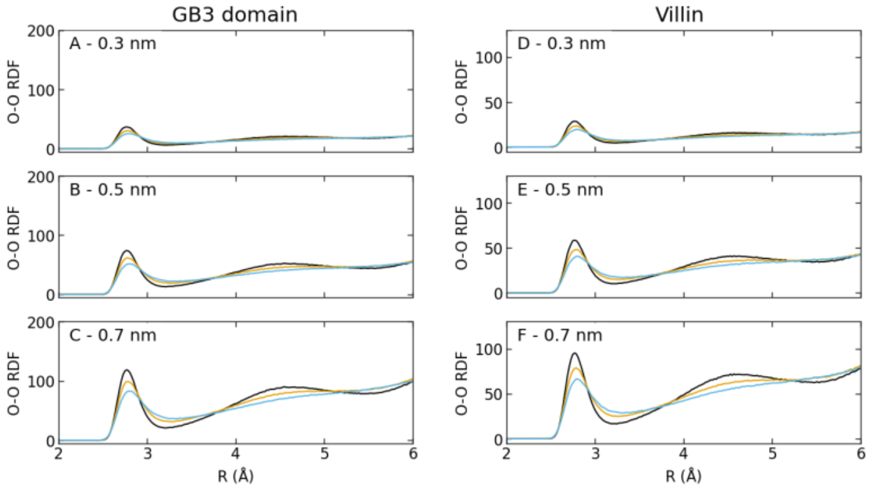


Figure S7: Analysis of the internal water structure within the hydration shell. (A) Non-normalized radial distribution functions (RDFs) between pairs of water oxygen atoms (O–O) for (A–C) the GB3 domain or (D–F) villin headpiece. RDFs are shown for temperatures of 250 K (black), 300 K (orange), and 350 K (blue). RDFs were computed for O–O pairs within a distances of (A/D) 0.3 nm, (B/E) 0.5 nm, or (C/F) 0.7 nm from the protein surface. Peaks of the RDFs decay with increasing temperature, indicating a loss of water structure within the hydration shell.
